# Cryo-FIB workflow for imaging brain tissue via *in situ* cryo-electron microscopy

**DOI:** 10.1101/2023.02.11.528064

**Authors:** Jiying Ning, Jill R. Glausier, Chyongere Hsieh, Thomas Schmelzer, Silas A. Buck, Jonathan Franks, Cheri M. Hampton, David A. Lewis, Michael Marko, Zachary Freyberg

**Affiliations:** Department of Psychiatry, University of Pittsburgh, Pittsburgh, PA 15213, USA; Biggs Laboratory Wadsworth Center, NYS Department of Health, Empire State Plaza, Albany, NY 12237, USA; TGS Technologies, Cranberry Township, PA 16066, USA; Center for Neuroscience, University of Pittsburgh, Pittsburgh, PA 15213, USA; Department of Cell Biology, University of Pittsburgh, Pittsburgh, PA 15261, USA; UES Inc. Dayton, OH 45433, USA; Materials and Manufacturing Directorate, Air Force Research Laboratory, Wright-Patterson Air Force Base, Dayton, OH, 45433, USA

## Abstract

Cryo-electron microscopy (cryo-EM) enables the study of protein complexes, cytoskeletal elements, and organelles in three dimensions without the use of chemical fixation. Most cryo-EM studies focus on vitreously frozen individual cells separated from their native tissue contexts. This reliance on imaging of single cells is primarily due to technical challenges associated with preparing fresh tissue sections at a thinness sufficient for visualization via cryo-EM. Highly heterogenous and specialized tissues, such as brain, are especially affected by this limitation as the cellular, subcellular, and synaptic milieus can significantly vary across neuroanatomical locations. To address this limitation, we established new instrumentation and a workflow that consists of: 1) high-pressure freezing of fresh brain tissue; 2) tissue trimming followed by cryo-focused ion beam milling via the H-bar approach to generate ultrathin lamellae; and 3) cryo-EM imaging. Here, we apply this workflow to visualize the fine ultrastructural details of organelles, as well as cytoskeletal and synaptic elements that comprise the cortical neuropil within fresh, unfixed mouse brain tissue. Moreover, we present initial studies that apply principles of the above workflow to the analysis of postmortem human brain tissue. Overall, our work integrates the strengths of cryo-electron microscopy and tissue-based approaches to produce a generalizable workflow capable of visualizing subcellular structures within complex tissue environments.

## Introduction

The cryo-electron microscopy (cryo-EM) revolution has enabled the visualization of macromolecules at near-atomic resolution without the need for chemical fixation. Indeed, cryo-EM single-particle analysis (SPA) is the ascendant approach for high-resolution structural reconstructions of macromolecules which are preserved in frozen-hydrated states via rapid vitrification (Lyumkis 2019, Wu and Lander 2020). However, most SPA samples are purified and therefore imaged *in vitro* outside of their native cellular environments. Therefore, the development of *in situ* cryo-imaging approaches that visualize structures in their native states within cells and in the absence of chemical fixation or other extrinsic manipulations that potentially distort cytoarchitecture promises many important biological insights (Oikonomou and Jensen 2017). The advent of *in situ* cryo-EM has begun delivering on this promise by imaging macromolecules and organelles directly within cells under near-native conditions including in healthy and disease states (Lucic, Leis et al. 2008, Oikonomou and Jensen 2017, Siegmund, Grassucci et al. 2018, Carter, Hampton et al. 2020).

To date, despite recent progress, most contemporary mammalian cryo-EM studies remain limited to individual cells grown in isolation. This runs contrary to mammalian physiology *in vivo* where cells function within the highly interactive tissue milieu. Indeed, mRNA and protein expression, as well as morphological differences exist between isolated cell types grown in culture versus cells within their native tissues (Hakkinen, Harunaga et al. 2011, Dauth, Maoz et al. 2017, Keil, Sethi et al. 2017, Lopes-Ramos, Paulson et al. 2017, Campiglio, Figliuzzi et al. 2021). Neurons are especially vulnerable to these limitations because they are highly differentiated and require inter-neuronal communication via synaptic contacts to function properly. Indeed, most neurons exhibit distinct, highly polarized cytoarchitectures that include morphologically complex pre-and postsynaptic processes that can span for millimeters or longer (Ho, Lee et al. 2011, Bailey, Kandel et al. 2015, Kulik, Watson et al. 2019). An individual neuron may receive upwards of 30,000 synapses, which represent the basic unit of neuronal communication (Megías, Emri et al. 2001, DeFelipe, Alonso-Nanclares et al. 2002, DeFelipe, Elston et al. 2002). These synaptic contacts are highly plastic, and make adaptive changes based on their local environments (Kandel 2001, Ho, Lee et al. 2011, Bailey, Kandel et al. 2015). As neuropsychiatric diseases are closely associated with disruptions of these complex synaptic connections (Jackson-Lewis, Jakowec et al. 1995, van Spronsen and Hoogenraad 2010, Marín 2012, Glausier and Lewis 2013, Marsden 2013, Glausier and Lewis 2018, Dhuriya and Sharma 2020), the ability to directly image neuronal sub-cellular compartments and connections in tissue contexts via cryo-EM can provide important new structural and functional insights into brain function and dysfunction. Indeed, cryo-EM studies have already begun to shed new light on the structural underpinnings of neuronal function including components of the cytoskeleton and vesicular machinery of synaptic neurotransmission (Fernández-Busnadiego, Zuber et al. 2010, Fernández-Busnadiego, Schrod et al. 2011, Martinez-Sanchez, Laugks et al. 2021).

Major technical challenges persist in applying cryo-EM to cells and even greater difficulties are associated with the cryo-imaging of tissues. Foremost, adherent mammalian cells are typically several microns thick, especially in the cell center, making it difficult to vitrify samples uniformly (Turk and Baumeister 2020). Furthermore, the thickness of intact cells embedded within vitreous ice is significantly greater than the inelastic mean-free path of 300 keV electrons (~300 nm). Thus, inadequate penetration of the electron beam prevents all but the thinnest parts of cells (~200-400 nm-thick) from being imaged. To improve penetration of the electron beam, cryo-focused ion beam (cryo-FIB) milling has been increasingly employed to generate lamellae sufficiently thin for cryo-EM without altering the vitreous state of the sample (Marko, Hsieh et al. 2006, Marko, Hsieh et al. 2007, Rigort, Bauerlein et al. 2012, Strunk, Wang et al. 2012). Given the substantially greater thickness of tissue specimens compared to individual cells grown in isolation, applications featuring tissue-based cryo-EM are much more limited, with even fewer studies specifically focusing on intact brain tissue (Hsieh, Schmelzer et al. 2014, He, Hsieh et al. 2017, Zhang, Zhang et al. 2021). This paucity of brain-based studies is based on several factors including challenging tissue preparation conditions since fresh, unfixed brain tissue is soft and difficult to handle. Though conventional EM preparations rely on strong chemical fixation and dehydration which improve tissue handling, these approaches can also alter ultrastructure and extracellular spaces within brain tissue (Schultz and Karlsson 1972, Korogod, Petersen et al. 2015, Tsang, Bushong et al. 2018). Vitreous freezing addresses these limitations by preserving brain tissue in a near-native state but possesses its own challenges. The high water content of brain tissue makes it difficult to freeze without addition of cryoprotectants (*e*.*g*., sucrose), which can introduce their own freezing artifacts (Rosene, Roy et al. 1986). The application of vitrified brain tissue to cryo-EM poses the additional difficulty of reliably obtaining sufficiently thin samples. The “lift-out” technique has attempted to address this challenge by enabling extraction of thin lamellae for cryo-EM following cryo-FIB-milling of tissue (Schaffer, Pfeffer et al. 2019). Yet, despite recent improvements, the lift-out approach remains technically challenging with extremely low success rates for most users, making it difficult to adapt to high-throughput applications (Schaffer, Pfeffer et al. 2019, Kuba, Mitchels et al. 2021).

To address these challenges, we have established a workflow for cryo-EM imaging of brain tissue (**Figure 1**). **(1)** We developed a methodology for high-pressure freezing (HPF) in a modified carrier that reliably vitrifies brain tissue. **(2)** Frozen brain tissue is exposed and trimmed via cryo-ultramicrotomy, followed by **(3)** cryo-FIB-milling of the trimmed tissue via the “H-bar” technique to efficiently generate ultrathin lamellae for **(4)** cryo-EM imaging of subcellular structures within their tissue contexts. Advantages of this workflow include utilizing the H-bar approach, which has a greater likelihood of successful cryo-transfer of milled lamellae of brain tissue and compatibility with modern cryo-microscopes. We also present initial studies that translate key principles of our workflow to the imaging of cryo-preserved postmortem human brain tissue.

**Figure 1.**
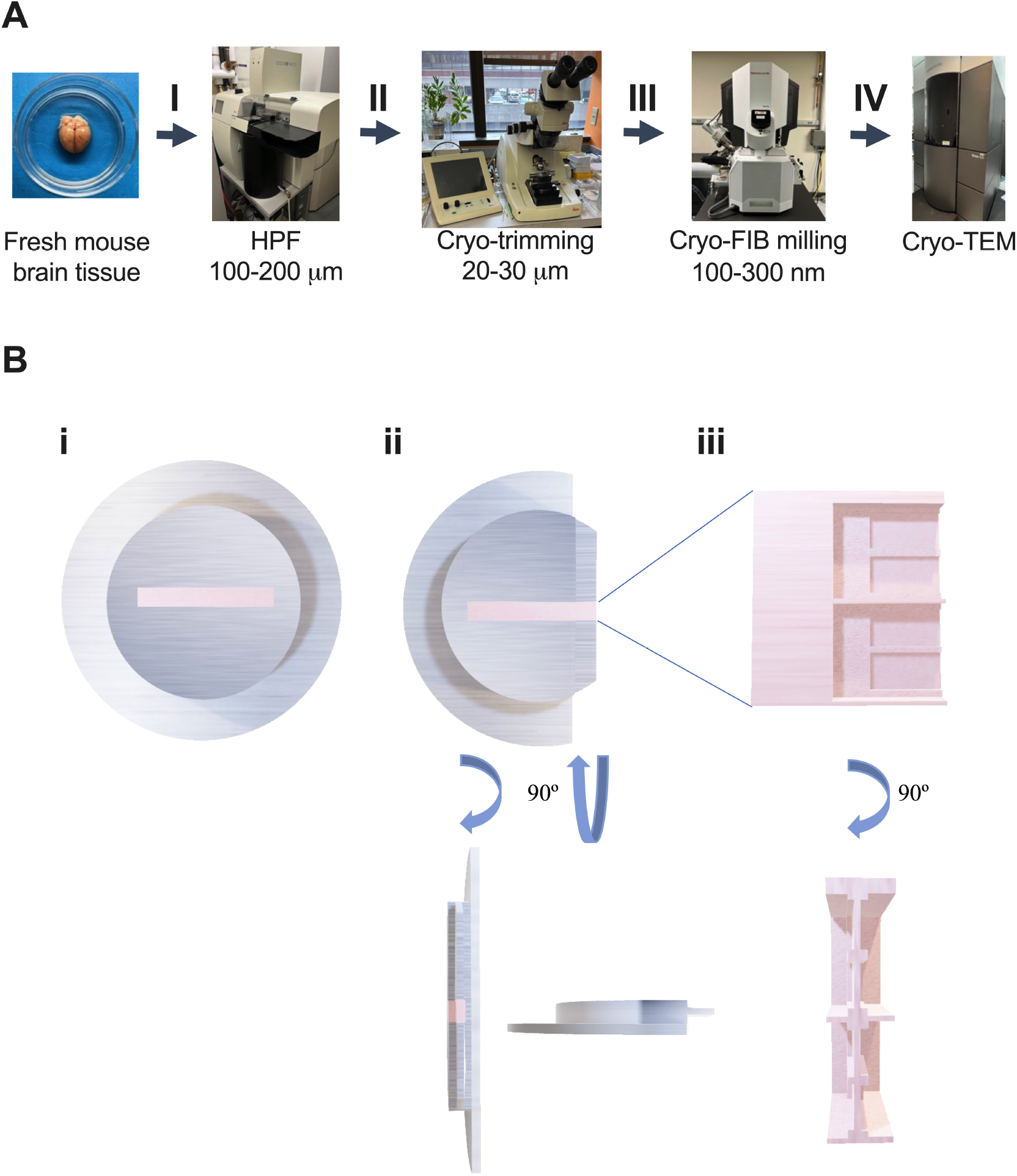
Workflow for preparation of fresh brain tissue for cryo-EM imaging. **(A)** Overview of the workflow. (**I**) 100-200 µm-thick sample of fresh mouse brain tissue is painted into an aluminum carrier. (**II**) Carrier is loaded into a high-pressure freezer (HPF) for immediate cryopreservation. (**III**) The HPF sample is subsequently trimmed in a cryo-ultramicrotome to a 20-30 µm thickness. (**IV**) The sample is thinned via cryo-FIB milling to a final 100-300 nm thickness using the H-bar approach, followed by imaging of the lamellae in a cryo-TEM. **(B)** Schematic of progressive thinning of the fresh brain sample in preparation for cryo-imaging. The HPF carrier (**Panel i**) is first trimmed away to expose brain tissue (**Panel ii** with 90° views). This is followed by progressive cryo-FIB-milling of lamellae as indicated in the inset (**Panel iii** with 90° views).

## Results and Discussion

We first established an approach to cryopreserve fresh brain tissue via HPF (see Methods). Tissue was dissected from fresh mouse frontal cortex. As such, brain tissue was not exposed to any fixation or contrast-enhancing agents that may distort ultrastructure. The tissue was placed into custom-made aluminum carriers using a fine-tipped brush, followed by immediate freezing via HPF.

We initially assessed the fresh, unfixed brain ultrastructure subsequent to HPF by processing a subset of these samples for freeze substitution, followed by imaging via conventional transmission electron microscopy (TEM). We identified neuronal compartments such as a dendritic spine connected to the parent dendritic shaft (**Figure 2A**), an axon terminal making a close apposition to a dendrite (**Figure 2B**), and organelles such as a large mitochondrion (**Figure 2C**). Thus, our manual approach to fresh brain tissue specimen collection prior to HPF sufficiently preserved ultrastructural quality of the grey matter neuropil.

**Figure 2.**
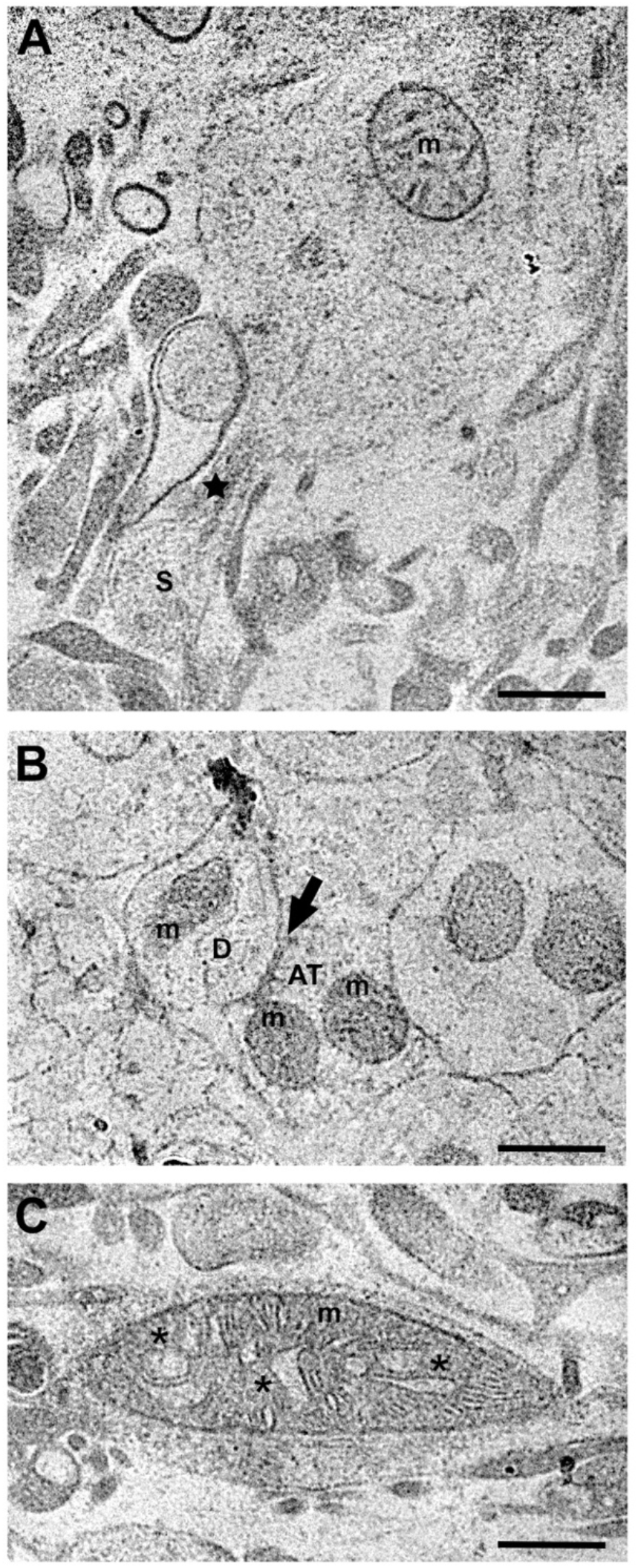
Ultrastructural preservation of fresh, unfixed mouse brain tissue via HPF/freeze substitution. **(A)** A dendritic spine (S) protruding from the parent pyramidal neuron dendritic shaft (D). The dendritic shaft contains a mitochondrion (m). A putative spine apparatus (star) is apparent in the spine neck. **(B)** A neuronal axon terminal (AT) containing two mitochondria (m) and forming an apposition (arrow) onto a dendritic shaft (D) containing a mitochondrion (m). **(C)** A large mitochondrion located within a dendritic shaft, displaying characteristics of a condensed state, which include an electron-dense matrix and multiple open cristae (*). Scale bars = 500 nm.

We next developed steps to prepare fresh, unfixed, HPF-preserved cortical tissue for cryo-EM. After HPF, the aluminum carriers containing cortical tissue were transferred to a holder termed the Intermediate Specimen Holder (ISH) which was custom-designed for this application (**Figure 3**). Upon mounting, the carrier was trimmed using a cryo-ultramicrotome where a portion of the aluminum casing was cut away to expose 20-30 µm-thick frozen-hydrated samples. This trimming step facilitated cryo-FIB-milling by restricting the amount of tissue required for subsequent thinning, significantly diminishing processing time and reducing risk of ice contamination. The pre-trimmed tissue was then cryo-FIB-milled via the H-bar approach to obtain 100-300 nm-thick lamellae which were subsequently visualized by cryo-EM.

**Figure 3.**
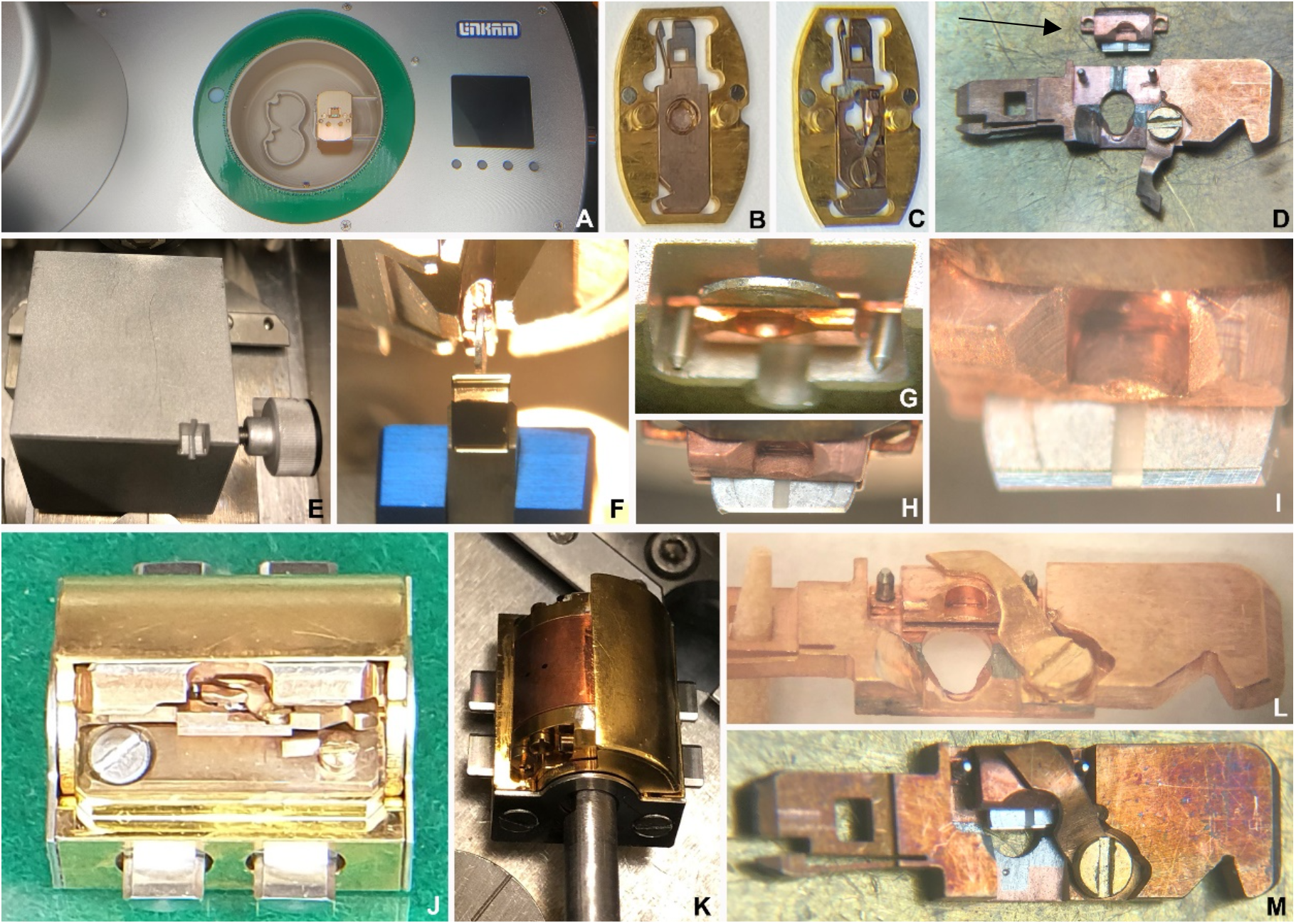
Devices for cryo-trimming and cryo-FIB milling of brain tissue. **(A)** Linkam CMS-196 cryo-microscope stage. In this view, a specially made frame for the Intermediate specimen holder (ISH) is shown. **(B)** A specially made frame for the cryo-stage holding an unmodified cartridge for the cryo-TEM. **(C)** The same frame, now holding the modified TEM cartridge with the ISH and HPF carrier mounted. **(D)** An ISH with a trimmed aluminum HPF carrier mounted within it (top, arrow), and a TEM cartridge placed alongside (bottom). **(E)** The loading block for the ISH. The jaws are slightly opened to accept the HPF carrier, which is then held (for cryo-FIB/SEM and cryo-TEM) by spring tension. **(F)** The ISH is held in the cryo-microtome chuck for the trimming step. **(G)** An untrimmed HPF carrier in the ISH. **(H)** After trimming, tissue is exposed. **(I)** Higher magnification image of the trimmed tissue in panel H. **(J)** The cryo-stage holder for the cartridge in the cryo-FIB, featuring an open shutter. **(K)** The cryo-stage holder is positioned on the rod to facilitate the cryo-transfer step from the cryo-FIB/SEM to the cryo-TEM. The shutter is shown closed, but automatically opens (or closes) when inserted into (or removed from) the platform of the FIB cryo-stage. **(L, M)** Two views of the cryo-TEM cartridge with the ISH in place.

Cryo-EM imaging of mouse brain revealed neuronal structures, organelles, and cytoskeletal elements (**Figure 4**). For example, we visualized neuronal axons containing microtubule tracks (**Figure 4A**). Within the microtubule lumen, we also identified intraluminal proteins, consistent with previously described work (Garvalov, Zuber et al. 2006, Foster, Ventura Santos et al. 2022). We also identified multiple mitochondria inside a neuronal process (**Figure 4B**). Within axon terminals, there were clusters of well-defined synaptic vesicles alongside a mitochondrion (**Figure 4C**). Finally, we identified an apposition between an axonal varicosity and a dendritic spine, which featured a spine apparatus (**Figure 4D**). Quantitative analyses revealed that mean neuronal plasma membrane thickness (4.4 ± 1.2 nm), synaptic vesicle diameter (41.0 ± 1.9 nm) and spacing of mitochondrial outer and inner membranes (18.5 ± 3.6 nm) were consistent with previously published values (Peters, Palay et al. 1991, Kühlbrandt 2015), demonstrating the preservation of neuronal ultrastructure in our cryo-preserved specimens.

**Figure 4.**
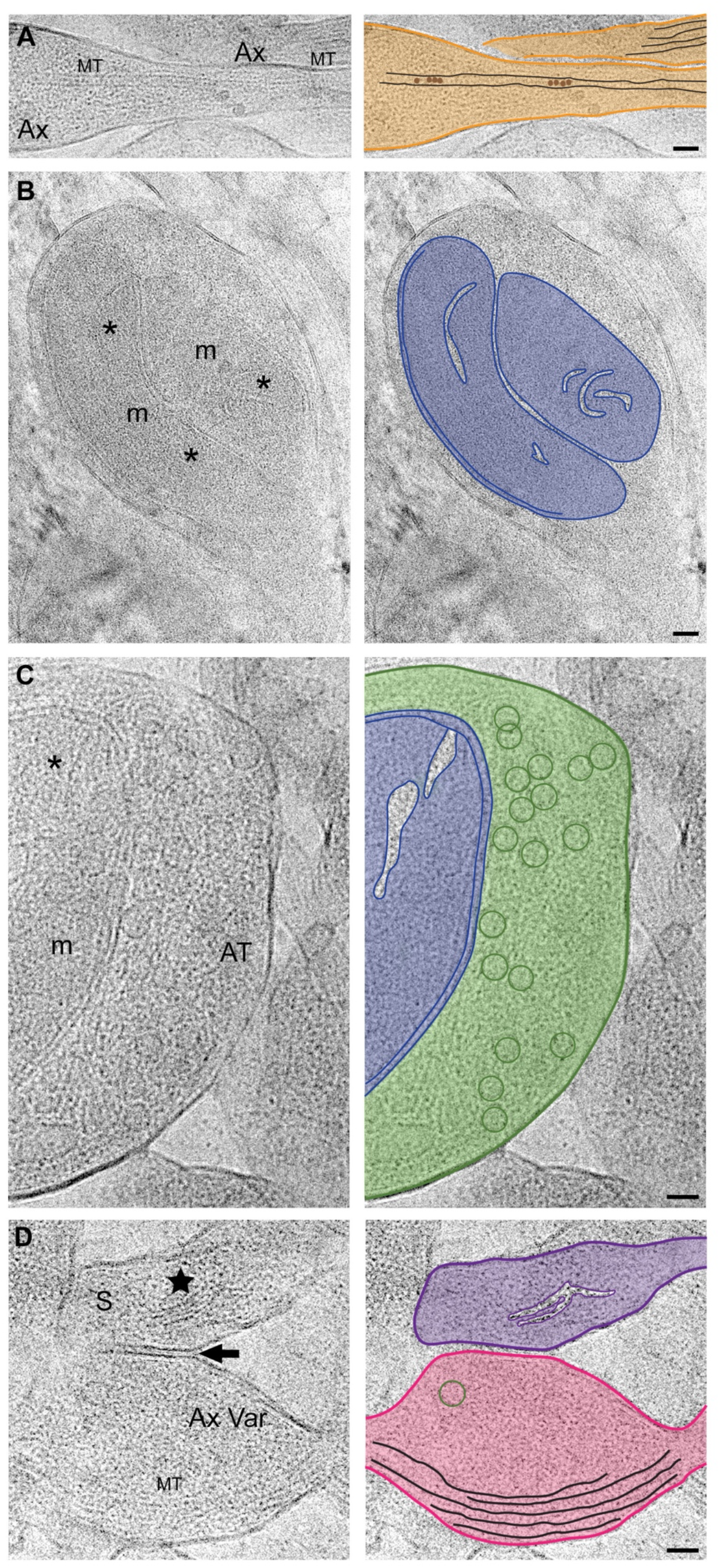
Representative 2D micrographs from cryo-FIB-milled lamellae of fresh, unfixed mouse cortical neuropil imaged via cryo-EM. Left panels are annotated original cryo-tomographic slices. Right panels feature segmentation of salient neuronal features. **(A)** Neuronal axons (Ax, orange) containing microtubule tracks (MT, black). Putative intraluminal proteins (brown) are visualized inside the central microtubule lumen. **(B)** Two nested mitochondria (m, blue) with visible cristae (asterisk) within a neuronal process. **(C)** Axon terminal (AT, green) containing clusters of synaptic vesicles (green circles) alongside a mitochondrion (m, blue) featuring open cristae (asterisk). **(D)** Apposition (arrow) between an axonal varicosity (Ax Var, pink) and a dendritic spine (S, purple), featuring a spine apparatus (star). The axonal varicosity contains a synaptic vesicle (green circle) and microtubule tracks (MT, black). Scale bars = 50 nm.

We next translated our HPF approach from mouse brain to human brain as the first step to establishing a workflow for postmortem human brain tissue. Here, we used lightly-fixed postmortem human cortical tissue for HPF cryo-preservation, followed by freeze substitution. TEM imaging revealed ultrastructural preservation of neuronal profiles, organelles, and synaptic contacts (**Figure 5**). We identified a neuronal nucleus which prominently featured a well-defined nuclear envelope replete with visible nuclear pores (**Figure 5A**). Multiple intracellular organelles were also evident including rough endoplasmic reticulum (RER, **Figure 5B**), ribosome-associated vesicles (a novel vesicular RER sub-compartment (Carter, Hampton et al. 2020), **Figure 5C**), and a mitochondrion (**Figure 5D**). We also found a myelinated axon ensheathed by five oligodendrocyte wraps (**Figure 5E**). Finally, we observed an asymmetric, Type 1 synapse formed by an axon terminal contacting a dendritic spine (**Figure 5F**). This successful HPF cryo-preservation sets the stage for future work in fresh, unfixed human brain tissue.

**Figure 5.**
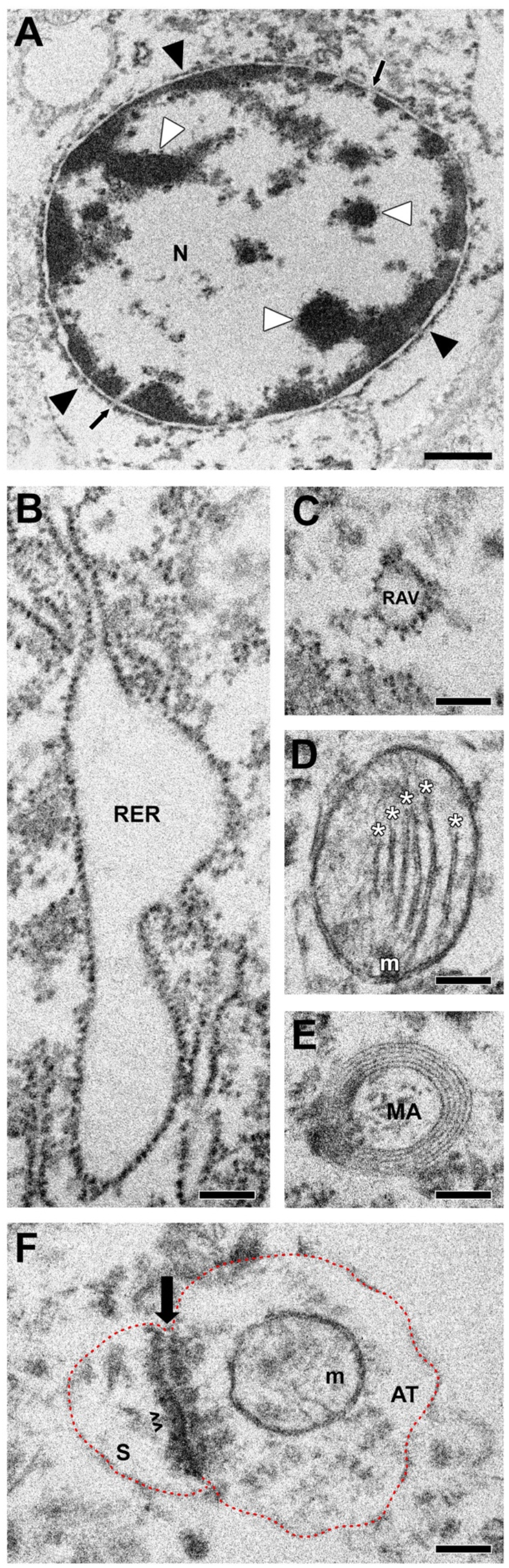
Ultrastructural preservation of lightly-fixed human brain tissue via HPF. **(A)** Neuronal nucleus (N) characterized by the presence of the nuclear envelope (black arrowheads) and chromatin aggregates (white arrowheads). The inner nuclear envelope membrane is smooth and lined internally by chromatin, whereas the outer membrane has a ruffled appearance. Multiple nuclear pores (arrows) are visible along the nuclear envelope. **(B)** Rough endoplasmic reticulum (RER) present within a neuronal cell body. **(C)** Ribosome-associated vesicle (RAV), a mobile form of RER, present within a neuronal cell body. **(D)** A mitochondrion exhibiting the characteristic double membrane enclosing the matrix and cristae (asterisk), which are regularly stacked. **(E)** Myelinated axon (MA) ensheathed by five regularly spaced oligodendrocyte wraps. **(F)** An asymmetric, Type 1 synapse formed by an axon terminal (AT) onto a dendritic spine (S). The synaptic contact features a separation of the axon terminal and spine by the synaptic cleft (arrow), as well as an electron-dense postsynaptic density (PSD) in the spine. Dotted line (in red) indicates putative plasma membrane outlining the axon terminal and the spine. Scale bar = 1 µm for **Panel A**. Scale bar = 200 nm for **Panels B-F**.

Future applications of this workflow will include *in situ* cryo-electron tomography (cryo-ET) imaging of mouse and postmortem human brain to obtain three-dimensional (3D) volumes of imaged structures. Additionally, the ability to apply sub-volume averaging methods to the data obtained from this approach will permit resolution of the 3D structures of target proteins in brain tissue. For example, recent work has explored the structures of pathogenic α-synuclein protein aggregates purified from brains of patients with Lewy pathology associated with Parkinson’s disease (Yang, Shi et al. 2022). The application of our workflow to resolve these disease-induced protein aggregates directly in human brain tissue of affected individuals will therefore represent a major advance in understanding disease pathogenesis. Finally, we also intend to incorporate cryo-correlative light and electron microscopy into this imaging workflow. The fluorescent signal would be correlated to specific sites within the brain sample to first guide sample trimming and then to guide subsequent cryo-FIB-milling to specific areas of interest expressing fluorescently labeled targets.

Overall, the above efforts to develop a workflow for *in situ* cryo-EM imaging of intact brain tissue promise to extend our understanding of cellular and subcellular interactions through visual mapping by cell-type and brain region. Furthermore, the addition of the H-bar approach for generating lamellae within tissue is less demanding and carries a higher likelihood of success compared to other approaches (*e*.*g*., lift-out). Our workflow also carries the eventual promise of sub-volume averaging which will enable direct 3D visualization of target proteins at sub-nanometer resolution in brain tissue. Such research approaches therefore have transformative potential to advance our understanding of basic neuroscience and drive the development of new treatments of brain diseases to improve human health.

## Materials and Methods

### Animals

C57BL/6J mice (The Jackson Laboratory, Bar Harbor, ME; JAX no. 000664) were maintained on a 12:12 hour light:dark cycle and with food and water available *ad libitum*. Animals were euthanized with isofluorane and transcardially perfused with PBS. Brains were rapidly removed and maintained in ice-cold PBS for the duration of tissue collection. All mouse experiments were approved by the University of Pittsburgh Institutional Animal Care and Use Committee (Protocol# 19075490). Animals were cared for in accordance with the ARRIVE guidelines for reporting animal research. All efforts were made to ameliorate animal suffering.

### Human Subject

The brain specimen was obtained during an autopsy conducted at the Allegheny County Office of the Medical Examiner (Pittsburgh, PA) after obtaining consent from next-of-kin. An independent committee of experienced research clinicians confirmed the absence of any lifetime psychiatric or neurologic diagnoses for the decedent based on medical records, neuropathology examinations, toxicology reports, as well as structured diagnostic interviews conducted with family members of the decedent (Glantz and Lewis 2000, Glausier, Kelly et al. 2020). The subject selected for study was a 45-year-old white male who died suddenly and out-of-hospital by natural causes from cardiovascular disease. The postmortem interval (defined as the time elapsed between death and brain tissue preservation) was 8.3 hours. Brain tissue pH was measured as 6.3, and RNA Integrity Number was determined as 8.5. These values reflect excellent tissue quality. All procedures were approved by the University of Pittsburgh’s Committee for the Oversight of Research and Clinical Training Involving Decedents and the Institutional Review Board for Biomedical Research.

### High-Pressure Freezing

For mouse studies, a semi-thin section of fresh, unfixed prefrontal cortex was taken manually using a #11 scalpel blade in the coronal plane and transferred to a petri dish filled with ice-cold PBS. Tissue from the superficial or middle cortical layers was further dissected, and then thinned using a fine-tipped brush (Dynasty, SC2157R, 5/0). The thinned tissue was transferred directly into custom-designed aluminum HPF carriers (Engineering Office M. Wohlwend GmbH, Sennwald, Switzerland). Carriers had a diameter of 3.0 mm, with a 0.5 mm recessed edge and a slot-shaped specimen well (1.4 mm ξ 0.2 mm ξ 0.15 mm) compatible with the Leica EMPACT2 HPF system (Leica Microsystems, Wetzlar, Germany) (Figure 1B). The carriers were manually loaded and torqued to 0.2 N-m in a flat specimen bayonet pod. HPF was immediately completed at a minimum pressure of 2000 bars with frozen samples stored in liquid nitrogen until further use.

For postmortem human brain studies, tissue was collected, fixed, and sectioned as previously described (Glausier, Konanur et al. 2019). Briefly, a small (~1 cm^3^) coronal block of dorsolateral prefrontal cortex was submerged in 4% paraformaldehyde/0.2% glutaraldehyde fixative for 24h at room temperature, followed by 24h at 4°C. The tissue block was then rinsed in 0.1M phosphate buffer (PB) and sectioned at 50 µm thickness via vibratome (VT 1000P, Leica). Tissue sections were stored in a cryoprotectant solution at −30°C until processed for HPF. Upon processing, sections were rinsed in ice-cold PB, and a sample of grey matter was transferred directly into a carrier. HPF, freeze substitution, ultrathin sectioning and lead staining were performed as described for mouse brain. Cryo-preserved tissue was examined using a JEOL 1400Plus TEM equipped with a 120 keV electron beam.

### Freeze Substitution for Screening

Initial evaluation of mouse and human brain specimen ultrastructure post-HPF was performed on samples processed via freeze substitution, as previously described (Yang, Bhatti et al. 2013). Briefly, frozen samples were warmed from −196°C to −90°C over three days in 1% OsO_4_/0.1% uranyl acetate (in acetone), gradually warmed to room temperature over 18h and subsequently embedded with EPON. Ultrathin sections were collected via ultramicrotome onto 200 copper mesh grids, lead-stained, and examined with a Tecnai T12 electron microscope (Thermo Fisher Scientific FEI).

### Cryo-Trimming

The carrier was mounted into the ISH, transferred into a UCT/FC cryo-ultramicrotome (Leica Microsystems). Using a diamond trimming knife (Diatome, Biel, Switzerland), a portion of the aluminum carrier was trimmed away to expose tissue. To reduce cryo-FIB-milling time, the sample was further trimmed to a 20-30 µm thickness. After trimming, the ISH-mounted carrier was stored under liquid nitrogen for later cryo-FIB-milling.

### Cryo-Transfer to the Cryo-FIB/SEM

The frozen, trimmed specimen was moved to the cryo-FIB/SEM for cryo-FIB-milling via cryo-transfer as described earlier (Hsieh, Schmelzer et al. 2014). Briefly, the ISH-mounted specimen was first loaded into a pre-tilted specimen block (Leica; **Figure 3 J&K**). The block was then placed into a Leica VCT-100 cryo-transfer system under liquid nitrogen to facilitate transfer into the pre-cooled cryo-FIB/SEM cryo-substage.

### Cryo-FIB-Milling

Using a Neon 40 EsB cryo-FIB/SEM (Zeiss, Oberkochen, Germany), the pre-trimmed ISH-mounted specimens were progressively thinned in a stepwise manner to reduce their thickness. Starting with an initial 20-30 µm thickness, samples were first milled to 10 µm, then to 3 µm, and finally to 100-300 nm thickness via the H-bar approach (Hsieh, Schmelzer et al. 2014, De Winter, Hsieh et al. 2021). The detailed steps of the H-bar method are described below:

Step 1: The thickness of the cryo-trimmed specimen was reduced to 10 µm in an area 200-250 µm wide by milling on both sides of the lamella (5 nA, 15 min) to generate a 60 µm-deep lamella.

Step 2: The thickness of the lamella was further reduced to 3 µm in two 80-µm-wide areas by cryo-FIB-milling (1 nA, 10 min) on both sides to create an “H” shape of ~40 µm depth.

Step 3: In the final cryo-FIB-milling step (100 pA, 3 min), the thickness of the lamella was reduced to ~100-300 nm within each of the above two areas, resulting in a lamella of sufficient thinness for cryo-EM imaging.

All cryo-FIB-milling steps were continuously monitored via SEM. Images were recorded with secondary electrons from the ion beam (30 keV, 10 pA) with frame integration and use of a ‘‘SESI’’ (Everhart–Thornley type) detector.

### Cryo-EM

Ultra-thin samples were loaded into a JEOL JEM-3200FSC/PP cryo-EM (JEOL Ltd., Tokyo, Japan) using a JEOL cartridge and cryo-transfer system; the cartridge was modified to fit the ISH-mounted specimen. Cryo-micrographs were recorded from the cryo-FIB-milled lamellae with a pixel size of 0.33 nm, 300 keV electron beam, zero-loss energy filtering (20 eV slit), 4 µm defocus, and electron dose of 100 e^-^/Å^2^.

### Ultrastructural Analysis

Neuronal profiles, organelles, and cytoskeletal elements were identified using established criteria summarized here (Peters, Palay et al. 1991). Neuronal nuclei were defined as round and containing both chromatin and a nuclear envelope. RER was located within somata and characterized by cisternae whose exterior surface was studded with electrondense ribosomes. RAVs were characterized by a ribosome-studded vesicular structure with no connections to a larger reticular ER network (Carter, Hampton et al. 2020). Microtubules were identified as linear structures ~20 nm in diameter and could contain intraluminal particles. Mitochondria were typically round or oblong in shape and identified by their characteristic double membranes encasing cristae and an electron-dense matrix. Axon terminals contained synaptic vesicles, could form synaptic contacts, and could contain mitochondria. Myelinated axons were defined by a central axonal structure surrounded by oligodendrocyte processes which form the characteristic myelin sheath. Dendritic shafts could contain microtubules and mitochondria, did not contain synaptic vesicles, and could receive synaptic contacts. Dendritic spines contained actin, could contain a spine apparatus, could receive synaptic contacts, and did not contain mitochondria, microtubules, or synaptic vesicles.

## Author contributions

Z.F. and M.M. initiated and supervised the project. J.N., J.R.G. performed HPF of mouse and human brain tissue with input from J.F. S.A.B. acquired fresh mouse brain tissue for the studies. J.R.G. and D.A.L. provided postmortem human brain tissue. J.N. and J.R.G. designed and tested the carriers. T.S. designed and machined the instrumentation specially adapted for the workflow. C.H. and M.M. performed cryo-trimming, cryo-FIB milling, and cryo-EM imaging of brain tissue. J.N., J.R.G., C.H., C.M.H., M.M., and Z.F. analyzed the data. J.N., J.R.G., M.M., and Z.F. wrote the manuscript with input from all the co-authors.

## Acknowledgments

This work was supported by the National Institutes of Health (R21DA052419, R21AA028800, R21AG068607 to Z.F.; R35GM119023 to M.M.), Commonwealth of Pennsylvania Formula Fund Award, the Pittsburgh Foundation (FPG00043-01 to Z.F.), as well as by the Department of Defense (PR192466, PR210207 to Z.F.). Postmortem human brain tissue was obtained from the NIH NeuroBioBank at the University of Pittsburgh. We are grateful to Dr. Donna Stolz and the Center for Biologic Imaging at the University of Pittsburgh for their ongoing support throughout this work. We thank Ming Sun for generous assistance in sample preparation. We also gratefully acknowledge the assistance of Mary L. Brady in the preparation of the figures.

